# Generation, Characterization and Comparison of Ovine Induced Pluripotent and Embryonic‑Disc Stem Cells

**DOI:** 10.64898/2026.04.30.721919

**Authors:** Catarina Silva-Almeida, Marta Esquiva Diaz, Waqas Ali, Sarah Ho, Maximilian Pickup, Sophia Webb, Deepika Rajesh, Patrick J Mee

**Affiliations:** Roslin Technologies, Roslin Innovation Centre, Easter Bush Campus, Edinburgh EH25 9RG, Scotland

**Keywords:** ovine, induced pluripotent stem cells, embryonic disc stem cells, differentiation, pluripotency, stability, transcriptomics, adipogenesis

## Abstract

Pluripotent stem cells derived from livestock species represent valuable systems for studying early mammalian development and for establishing renewable, well-defined cell sources; however, direct comparative characterization of distinct pluripotent stem cell platforms in sheep remains limited. In this study, we established and evaluated two ovine pluripotent stem cell types: reprogrammed induced pluripotent stem cells (siPSCs) and embryonic disc-derived stem cells (sEDSCs). Both siPSCs and sEDSCs exhibited core features of pluripotency, including compact colony morphology, alkaline phosphatase activity, expression of key pluripotency-associated markers, and maintenance of a normal ovine karyotype. Flow cytometry and quantitative RT-PCR analyses revealed broadly overlapping yet distinguishable pluripotency marker expression profiles between the two cell types. Functional pluripotency was confirmed by embryoid body formation and *in vitro* differentiation into derivatives of all three germ layers. To further assess lineage-specific differentiation competence and compare functional outputs relevant to mesodermal differentiation, both pluripotent stem cell types were directed towards the adipogenic lineage. While siPSCs and sEDSCs were each capable of adipogenic differentiation, differences in differentiation efficiency and marker expression were observed. Together, these findings demonstrate that ovine siPSCs and sEDSCs share core pluripotency characteristics while retaining distinct molecular and functional properties, providing a robust comparative framework for studies of ovine pluripotency, lineage specification, and stem cell biology.

## Introduction

Pluripotent stem cells (PSCs) provide powerful experimental systems for investigating early mammalian development and for enabling genetic, cellular, and translational technologies across species. Extensive work across mouse, human, and livestock systems has refined the molecular definition of pluripotency, identified multiple developmental pluripotent states, and clarified the signaling and epigenetic mechanisms that sustain these states *in vitro* (1–3). Together, this establishes pluripotent stem cells as *in vitro* surrogates for embryonic epiblast lineages and as powerful platforms for comparative studies of mammalian development (4,5). In addition, PSCs are increasingly being explored as renewable, well-defined cell sources for cellular agriculture, including cultivated meat production. In this context, pluripotent cells offer attractive properties, including long-term self-renewal and the potential to generate diverse lineages (6–8). Recent progress in stem cell-based approaches for cultivated meat, including demonstrations of serum-free differentiation and multi-lineage outputs from stable livestock pluripotent cell states, further supports the feasibility of PSC platforms for food relevant applications (9,10).

Early efforts to establish pluripotent stem cell lines in sheep, including both embryo-derived stem cells and reprogrammed induced pluripotent stem cells, were frequently characterized by limited stability, continued dependence on exogenous factor expression, or uncertainty regarding developmental identity (11). These challenges reflected, in part, an incomplete understanding of pluripotency regulation in the ovine embryo, as many derivation strategies were adapted from rodent or human systems without accounting for species-specific differences.

More recently, transcriptomic analyses of ovine pre-implantation embryos have revealed that pluripotency emerges through a dynamic progression of epiblast states, governed by distinct gene regulatory networks and shifting signaling dependencies during early development (5). These findings highlighted that ovine pluripotency does not conform cleanly to classical naïve or primed definitions established in other species, providing a biological explanation for the instability and ambiguity observed in earlier sheep pluripotent stem cell lines.

Against this backdrop, the generation of ovine induced pluripotent stem cells (iPSCs) offered an alternative route to pluripotency capture that bypasses direct derivation from embryonic tissue. Initial studies demonstrated that sheep fibroblasts could be reprogrammed into iPSC-like cells using inducible expression of defined transcription factors (12). These early ovine iPSCs expressed core pluripotency markers, formed embryoid bodies, and differentiated into derivatives of all three germ layers *in vitro*, establishing proof-of-principle for somatic cell reprogramming in sheep. Subsequent studies explored a range of somatic starting populations, factor combinations, and delivery systems, including viral, episomal, and transposon-based approaches, reflecting ongoing efforts to optimize ovine iPSC derivation and stabilization (13,14).

More recent ovine iPSC studies have increasingly employed piggyBac-based or doxycycline-inducible systems incorporating extended reprogramming factor sets. These approaches have reported stable colony morphology, expression of pluripotency-associated genes, karyotypic normality, and trilineage differentiation capacity, supported in some cases by transcriptomic profiling (15) Building on this work, additional studies have applied RNA sequencing to investigate signaling and metabolic pathways associated with pluripotency maintenance in ovine iPSCs, including MAPK, PI3K–AKT, TGF-β, and WNT signaling networks (16) Ovine iPSCs have also been assessed in developmental contexts, such as somatic cell nuclear transfer assays and early embryo development models, highlighting both their potential utility and ongoing challenges in defining fully reprogrammed pluripotency states (4)

In parallel with iPSC-based approaches, significant progress has been made in the derivation of embryo-derived pluripotent stem cells from livestock embryos under defined conditions. Notably, stable pluripotent embryonic stem cell lines have been derived from bovine blastocysts exhibiting sustained self-renewal, stable pluripotency marker expression, transcriptomic coherence, and competency in somatic cell nuclear transfer assays (17). Building on these advances, pluripotent stem cell lines were derived from porcine, ovine, and bovine embryos using feeder-free culture conditions incorporating activin A, fibroblast growth factor, and WNT inhibition. Global transcriptomic analyses placed these cells in close proximity to the bilaminar embryonic disc epiblast rather than the pre-implantation inner cell mass (2,18).This work distinguished these livestock embryo-derived stem cells from canonical mouse embryonic stem cells. Complementary ovine studies have independently reported the derivation of pluripotent stem cells from sheep blastocysts under defined signaling conditions, reinforcing the conclusion that ovine pluripotency can be stabilized *in vitro* but requires species-specific culture optimization (17,19,20)

Ovine iPSCs offer flexibility in donor selection and accessibility of starting materials but can exhibit variability in stability, continued dependence on exogenous factors, and heterogeneity in pluripotency state. Conversely, embryo-derived stem cells provide a biologically grounded reference for ovine pluripotency but require careful characterization to define their long-term behavior and differentiation potential. Despite a growing number of reports describing ovine iPSCs or embryo-derived pluripotent stem cells independently, there has been limited direct comparison of these platforms using aligned derivation and characterization strategies.

In this study, we established and systematically compared ovine siPSCs and sEDSCs using a unified experimental framework incorporating a common set of orthogonal assays to assess pluripotency, genomic stability, transcriptional state, and differentiation potential. This strategy enables a structured evaluation of reprogrammed and embryo-derived pluripotent stem cell platforms in sheep and clarifies their respective properties as *in vitro* models of ovine pluripotency.

## Materials and Methods

### Derivation of ovine embryonic disc stem cells

Frozen sheep embryos (ABreeds) were received in sealed straws and thawed according to the supplier’s instructions. Following thawing, embryos were flushed from each straw using pre-equilibrated BO IVC medium (IVF Bioscience, Cat. No. 7001) and examined under an inverted microscope to assess developmental stage, morphology, and the presence of an intact zona pellucida. Blastocysts retaining an intact zona pellucida were subjected to chemical dezoning using Acid Tyrode’s solution (Sigma-Aldrich, Cat. No. T1788) until complete zona removal was confirmed microscopically. All intermediate handling and washing steps were performed using Dulbecco’s phosphate-buffered saline (DPBS) (Gibco, Cat. No. 14190-144) supplemented with 0.2% (w/v) polyvinylpyrrolidone (PVP) (Sigma-Aldrich, Cat. No. P0930).

Zona-free blastocysts were processed by immunosurgery to isolate the inner cell mass (ICM). Embryos were incubated in 20% (v/v) sheep serum antibody solution (Rockland, Ct. No. 113-4101) for 1 h at 37 °C, with intermittent washing in PVP-supplemented DPBS to remove unbound antibodies. Embryos were subsequently transferred to 20% (v/v) guinea pig complement (Sigma-Aldrich, Cat. No. S1639) and incubated for a further 45 min at 37 °C. Following complement-mediated lysis, trophectodermal cells were removed by gentle repeated pipetting, and intact ICMs were visually identified and collected.

Individual ICMs were plated into single wells of 12-well tissue-culture plates coated with fibronectin (Merck, Cat. No. ECM001) and laminin-511 (Merck, Cat. No. CC160) prepared according to the manufacturers’ instructions. ICMs were cultured as previously described (18) however ICMs were initially cultured in JXGO media (Roslin Technologies) supplemented with activin A prior to culture in AFX medium (DMEM/F-12 backbone and combines fibroblast growth factor and activin/TGF-β signaling with inhibition of WNT pathway activity). Cultures were left undisturbed for the first 48 h to allow ICM attachment, after which medium was replaced daily. Emerging outgrowths were manually dissected and expanded by passaging onto freshly coated wells under identical culture conditions.

### Primary cell lines

Neural progenitor cells were derived from the subventricular zone of the brain as previously described (21). Briefly, brain tissue was micro-dissected, enzymatically dissociated, and plated onto tissue-culture–treated 6-well plates (Corning Costar, Cat. No. 3516) that had been coated with laminin-511 (Merck, Cat. No. CC160) according to the manufacturer’s instructions. Cells were maintained in neural stem cell medium RHB-A (Takara Bio, Cat. No. Y40001) with 9 µg/mL recombinant human epidermal growth factor (EGF) (PeproTech, Cat. No. AF-100-15), and 9 µg/mL recombinant human basic fibroblast growth factor (FGF-2) (PeproTech, Cat. No. 100-18B). Neural cultures were maintained under standard humidified conditions at 37 °C and 5% CO_2_ until adherent neural populations emerged, typically within 8–14 days. Cells were subsequently passaged using TrypLE™ Express Enzyme (Gibco, Cat. No. 12604013) and expanded for downstream applications.

Primary fibroblasts were derived from skin biopsies using an explant outgrowth method. Briefly, skin samples were minced into approximately 1–2 mm^2^ explants and placed into uncoated tissue-culture plates containing fibroblast growth medium composed of DMEM/F-12 (Gibco, Cat. No. 11320-033) supplemented with 10% fetal bovine serum (FBS) (Avantor, Cat. No. 97068-085), 1× NEAA (Gibco, Cat. No. 11140050), 1× sodium pyruvate (Gibco, Cat. No. 11360070), and 0.1 mM 2-mercaptoethanol (Gibco, Cat. No. 21985023). Explants were maintained until fibroblast outgrowth was observed, after which cells were passaged using TrypLE™ Express and expanded.

All ovine tissues were obtained via a licensed abattoir. All media used for primary cell derivation were supplemented with Antibiotic–Antimycotic (100×) solution (Sigma-Aldrich, Cat. No. A5955) and Primocin™ (InvivoGen, Cat. No. ant-pm-1) to minimize microbial contamination during primary culture establishment.

### Reprogramming and culture of ovine induced pluripotent stem cells

Primary ovine neural cells were seeded onto culture plates coated with laminin-511 (Merck, Cat. No. CC160) and allowed to attach for 24 h prior to reprogramming. Cell numbers were determined to calculate viral input, after which cells were transduced using the CytoTune™-iPS 2.0 Sendai Reprogramming Kit (Thermo Fisher Scientific, Cat. No. A16517) expressing the four Yamanaka factors OCT3/4, SOX2, KLF4, and c-MYC. Viral infectivity was confirmed using the CytoTune™ EmGFP Sendai Fluorescence Reporter (Thermo Fisher Scientific, Cat. No. A16519).

Twenty-four hours post-transduction, culture medium was replaced, and cells were maintained for 6– 7 days in neural stem cell medium as described above. Seven days after transduction, cells were dissociated using TrypLE™ Express Enzyme and replated onto laminin-511-coated plates in activin-containing pluripotent stem cell medium. Emerging colonies with characteristic pluripotent morphology were manually picked and transferred into laminin-511-coated 24-well plates for expansion.

Established siPSCs were maintained in JXGO medium (Roslin Technologies) under feeder-free conditions, passaged every 6–8 days using TrypLE™ Express, and replated at a density of 7.3 × 10^3^ cells/cm^2^ onto laminin-511-coated culture vessels. Cultures were maintained at 37 °C in a humidified atmosphere of 5% CO_2_.

### Alkaline phosphatase stain

Alkaline phosphatase (AP) activity was assessed in emerging stem cell colonies as a marker of maintenance of an undifferentiated state. Cells were cultured until near confluence, washed with Dulbecco’s phosphate-buffered saline (DPBS) (Gibco, Cat. No. 14190-144), and fixed with 4% paraformaldehyde (Thermo Scientific, Cat. No. J61889.AK) for 10 min at room temperature. Following fixation, cells were washed with DPBS and stained using the AP Detection Kit (SLS, Cat. No. 86R) according to the manufacturer’s instructions. Briefly, staining solution was applied to fixed cultures and incubated for 30 min at room temperature, after which plates were washed with DPBS and imaged by bright-field microscopy.

### Quantitative RT-PCR

Total RNA was isolated from ovine siPSCs and ovine sEDSCs using the RNeasy Mini Kit (Qiagen, Cat. No. 74104) according to the manufacturer’s instructions. RNA concentration and purity were assessed spectrophotometrically. Complementary DNA (cDNA) was synthesised from 500 ng total RNA using the AffinityScript cDNA Synthesis Kit (Agilent Technologies, Cat. No. 600559) following the supplier’s protocol.

Quantitative PCR (qPCR) was performed using a QuantStudio™ 3 Real-Time PCR System (Applied Biosystems). Reactions were carried out using SYBR Green-based chemistry, with each 10 µL reaction containing 5 µL 2× SYBR Green PCR Master Mix (Applied Biosystems, Cat. No. A25742), 0.4 µL primer mix (5 µM each forward and reverse primer), 3.6 µL nuclease-free water, and 1 µL cDNA template (10 ng/µL). All reactions were prepared in technical triplicate using 96-well optical PCR plates, and no-template and no-reverse-transcription controls were included for each primer pair.

Thermal cycling conditions were as follows: initial incubation at 50 °C for 2 min, polymerase activation at 95 °C for 10 min, followed by 40 amplification cycles of 95 °C for 15 s and 60 °C for 1 min. Melt-curve analysis was performed to confirm amplification specificity, consisting of 95 °C for 1 s, 60 °C for 20 s, and 95 °C for 1 s with continuous fluorescence acquisition.

Ct values were determined using QuantStudio™ Design and Analysis Software v2.6.0 (Applied Biosystems). Relative gene expression was calculated using the ΔΔCt method. Expression levels were normalized to the housekeeping gene HMBS, and data were expressed relative to sheep fibroblasts (sFibs) as the calibrator sample. Primer sequences are listed in Table 1.

**Table 1.**
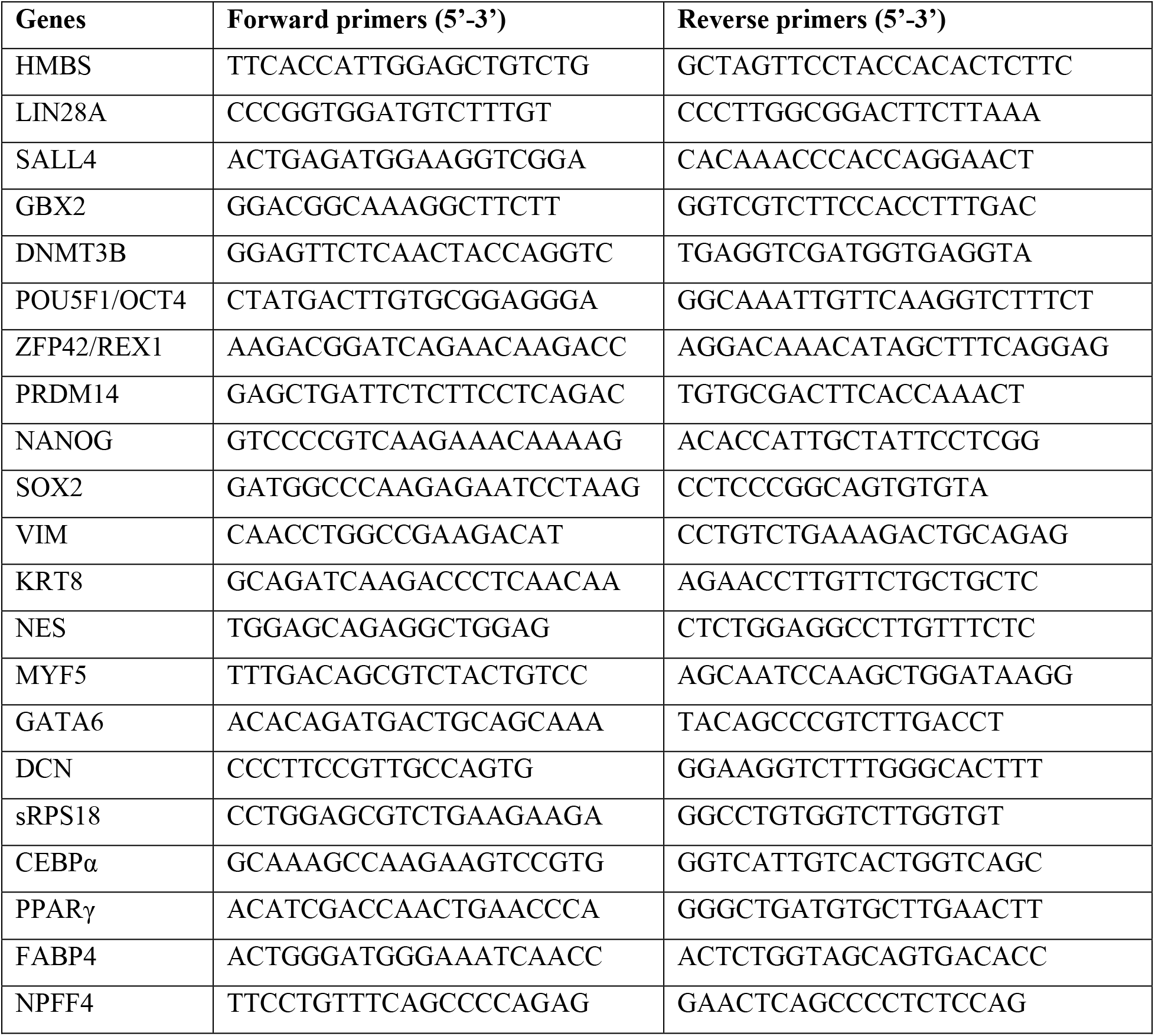
Primer sequences used for quantitative RT-PCR analysis of pluripotency-associated, lineage-specific, and housekeeping genes. All primers were designed to amplify ovine transcripts and were validated for specificity prior to use. Forward and reverse primer sequences are shown in the 5′–3′ orientation.

### Karyotyping

Karyotype analysis was performed by G-banding to assess chromosomal integrity. Cytogenetic analysis was conducted by Creative Bioarray (USA) or Cell Guidance Systems (UK), with a minimum of 20 G-banded metaphase spreads analyzed per sample. For metaphase arrest, cells were cultured to near confluence and incubated with KaryoMAX™ Colcemid™ Solution (Thermo Fisher Scientific, Gibco™, Cat. No. 15212012) for 1 h at 37 °C. Cells were dissociated using TrypLE™ Express Enzyme and collected into centrifuge tubes containing fetal bovine serum to quench enzymatic activity.

Following centrifugation, cell pellets were resuspended dropwise in pre-warmed 0.075 M potassium chloride hypotonic solution (Invitrogen, Cat. No. AM9640G) and incubated at 37 °C for 20 min to promote cellular swelling. Cells were then centrifuged and fixed using a freshly prepared 3:1 (v/v) methanol:acetic acid fixative. Fixed cell suspensions were stored at 4 °C prior to shipment for G-banding and cytogenetic evaluation.

### Embryoid body formation and in vitro differentiation

Embryoid bodies (EBs) formation was used to assess *in vitro* differentiation capacity. Ovine induced pluripotent stem cells (siPSCs) cultured to near confluence were dissociated into single cells using TrypLE™ Express Enzyme, centrifuged, and resuspended in EB differentiation medium at a density of 6,000 cells per EB in a total volume of 200 µL per well. EB medium was adapted from Liu et al. (2021) and consisted of KnockOut™ DMEM (Gibco™, Cat. No. 10829018) supplemented with 10% fetal bovine serum, 1× non-essential amino acids (NEAA; Gibco™, Cat. No. 11140050), and 1× GlutaMAX™ (Gibco™, Cat. No. 35050061).

Cell suspensions were seeded into ultra-low-attachment 96-well U-bottom plates (Corning®, Cat. No. 7007), with one EB formed per well. After 3 days in suspension culture, EBs were transferred to 6-well tissue culture plates pre-coated with 0.2% (w/v) gelatin (Sigma-Aldrich, Cat. No. G1890) to allow adherence and further differentiation. Cultures were maintained for up to 13 days, with medium changes performed every other day. At the end of the differentiation period, cells were harvested for RNA isolation and quantitative RT-PCR analysis of germ layer-specific marker expression.

The same protocol was used for EB generation from ovine sEDSCs, with the addition of 50 µM ROCK inhibitor Y-27632 (Enzo Life Sciences, Cat. No. ALX-270-333-M005) during the initial seeding on day 0 to enhance cell survival.

### Adipogenic differentiation

Adipogenic differentiation was performed using an adaptation of previously described human adipogenic differentiation protocols (22). Ovine pluripotent stem cells were cultured under standard maintenance conditions until near confluence prior to induction of adipogenesis. Differentiation was initiated using an induction phase incorporating all-trans retinoic acid to promote mesodermal lineage commitment, followed by exposure to the defined adipogenic differentiation cocktail containing insulin, dexamethasone, 3-isobutyl-1-methylxanthine (IBMX), the PPARγ agonist rosiglitazone, and triiodothyronine (T3) to support adipogenic maturation. Basal medium composition was adjusted to account for platform-specific differences between induced pluripotent stem cells (siPSCs) and embryonic disc-derived stem cells (sEDSCs). Cultures were maintained with regular medium changes until day 16, after which adipogenic conversion was assessed by Oil Red O staining (22) and quantitative RT-PCR analysis of adipocyte-associated gene expression.

### Flow cytometry

Flow cytometry was performed to quantitatively assess the expression of surface and intracellular pluripotency-associated markers in siPSCs and sEDSCs and to enable comparative phenotypic analysis between pluripotent stem cell platforms. Cells were dissociated into single-cell suspensions using TrypLE™ for 15 min at 37 °C, washed with DPBS, and resuspended in flow cytometry staining buffer (Cell Staining Buffer, BioLegend, Cat. No. 420201). Cell viability was assessed using LIVE/DEAD™ Fixable viability dyes (Invitrogen™), specifically the LIVE/DEAD™ Fixable Green Dead Cell Stain Kit (Cat. No. L23101) or LIVE/DEAD™ Fixable Violet Dead Cell Stain Kit (Cat. No. L34955), applied for 15 min at room temperature in the dark according to the manufacturers’ instructions.

For surface marker analysis, cells were incubated with fluorochrome-conjugated antibodies recognizing the pluripotency-associated surface antigens SSEA-1 or SSEA-4 for 30 min at room temperature in the dark. Following staining, cells were washed and fixed with 4% paraformaldehyde.

For intracellular marker analysis, cells were fixed with 4% paraformaldehyde and permeabilised for 20 min using 0.1% Triton X-100 (Sigma-Aldrich, Cat. No. X100) prepared in DPBS. Cells were subsequently stained for the pluripotency-associated transcription factors POU5F1 (OCT4), SOX2, LIN28A, or NANOG, using commercially available primary antibodies, followed where required by appropriate fluorophore-conjugated secondary antibodies (12). All antibody incubations were performed for 1 h at room temperature in the dark. Isotype-matched controls were included for all analyses.

Data acquisition was carried out using a BD LSRFortessa™ X-20 flow cytometer (BD Biosciences). Flow cytometry data were analyzed using FlowJo™ software (BD Biosciences), with appropriate compensation, viability gating, and exclusion of doublets applied during analysis.

### RNA sequencing Analysis

siPSCs, sEDSCs, primary neural cells, and fibroblasts were cultured to near confluence, and total RNA was extracted using the RNeasy Mini Kit (Qiagen, Cat. No. 74104) according to the manufacturer’s instructions. Library preparation and sequencing were performed externally by GENEWIZ (Azenta Life Sciences) using TruSeq Stranded mRNA chemistry (Illumina TruSeq Stranded mRNA Library Prep Kit, Cat. Nos. RS-122-2101 and/or RS-122-2102, depending on index set). Libraries were sequenced on an Illumina NovaSeq 6000 platform to generate 150 bp paired-end reads, targeting ∼30 million reads per sample.

Raw FASTQ files were assessed for quality using FastQC (v0.12.1). Adapter sequences and low-quality bases were trimmed using Cutadapt (v3.2) (23), removing reads containing adapter contamination, reads with Phred quality <30, reads <50 bp after trimming, and reads containing ambiguous bases (N) at the 5′ or 3′ ends. Trimmed reads were aligned to the Ovis aries reference genome NCBI ARS-UI_Ramb2.0 (GCF_016772045.1) using STAR (v2.7.11a) (24). Gene-level read counts were generated using featureCounts (v2.0.6) (25). Differential gene expression analysis was performed in R using edgeR (v4.2.2) (26). Genes were considered differentially expressed at FDR < 0.05 with an absolute log2 fold-change ≥ 1.5. Normalized counts were used to generate heatmaps for gene expression level using pheatmap package (v1.0.12) and differentially expressed genes were plot as a volcano plot using EnhancedVolcano package (v1.14.0).

## Results

### Establishment of ovine embryonic disc-derived and induced pluripotent stem cell lines

To enable a controlled comparison of ovine pluripotent stem cell platforms in subsequent experiments, embryonic disc-derived stem cells and induced pluripotent stem cells were first established as independent well characterized reference cell banks within the same study.

Sheep embryonic disc-derived stem cells (sEDSCs) were established from blastocyst-stage embryos (approximately day 6–7 post-conception). Following thawing and removal of the zona pellucida, embryonic material containing the inner cell mass was plated under feeder -free conditions incorporated FGF-mediated MAPK signaling, activin/TGF-β pathway activation, and LIF-associated STAT3 signaling, together with small-molecule modulation of WNT and protein kinase C (PKC) pathways (18) in normoxic conditions. Cultures gave rise to colony formation within two weeks of culture with sEDSC forming compact colonies with smooth, well-defined borders and minimal spontaneous differentiation (Figure 1). This colony morphology was maintained during serial passaging and subsequent cell banking.

**Figure 1.**
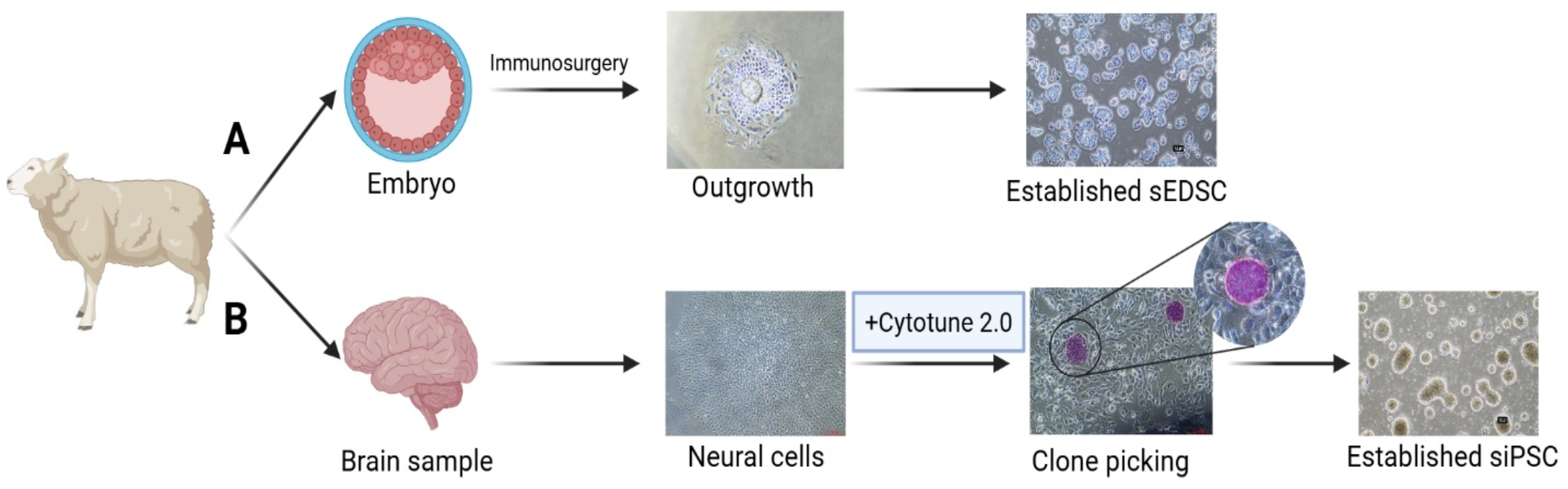
Overview of the derivation workflows and representative images illustrating the establishment of ovine pluripotent stem cell lines. (A) Sheep embryonic disc-derived stem cells (sEDSCs) were generated from blastocyst-stage embryos following immunosurgery. Representative phase-contrast images show initial embryonic outgrowths and the establishment of compact sEDSC colonies following expansion *in vitro*. (B) Sheep induced pluripotent stem cells (siPSCs) were generated from primary neural cells isolated from postnatal ovine brain tissue. Neural cells were expanded and reprogrammed using a non-integrating Sendai virus-based system (CytoTune™ 2.0), followed by manual selection and expansion of reprogrammed colonies. Representative phase-contrast images illustrate the neural starting population, emerging reprogrammed clones during colony picking, and established siPSC colonies. Representative alkaline phosphatase staining demonstrates robust AP activity in established sEDSC and siPSC cultures, consistent with a pluripotent stem cell phenotype. Scale bars as indicated.

Sheep induced pluripotent stem cells (siPSCs) were generated from primary neural cells isolated from postnatal ovine tissue, selected as a somatic starting population due to their proliferative capacity and permissive epigenetic state for reprogramming. Neural cultures were expanded under progenitor-supportive conditions prior to induction of pluripotency using a non-integrating Sendai virus–based system encoding the Yamanaka transcription factors OCT4, SOX2, KLF4, and c-MYC (27) under defined feeder-free media conditions that support pluripotency through TGF-β/activin signaling and FGF-dependent pathways (Figure 1).

### Shared pluripotency-associated feature

Following establishment of ovine sEDSC and siPSC we performed concurrent characterization of pluripotency-associated features using complementary morphological, molecular, and cytogenetic assays. Both siPSCs and sEDSCs formed compact, well-defined colonies and stained positively for AP, consistent with maintenance of an undifferentiated pluripotent state (Figure 2A). While overall colony morphology was similar, siPSC colonies exhibited a more compact, dome-shaped, three-dimensional architecture, whereas sEDSC colonies typically appeared flatter and more epithelial, reflecting differences observed during derivation (Figure 2A).

**Figure 2.**
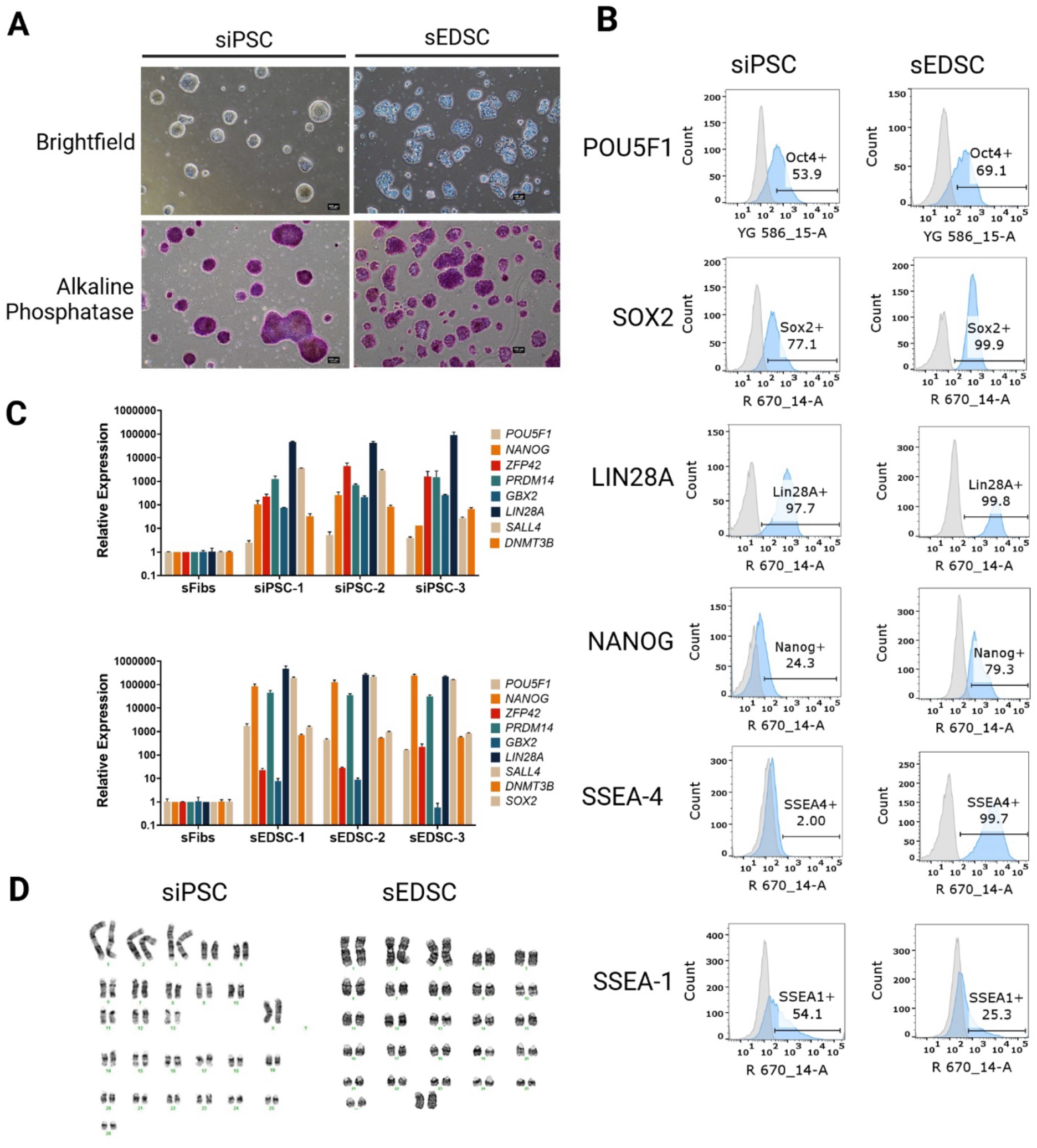
Morphological, molecular, and chromosomal characterization of ovine induced pluripotent stem cells (siPSCs) and embryonic disc-derived stem cells (sEDSCs). (A) Representative phase-contrast images (top row) and alkaline phosphatase staining (bottom row) of established siPSC and sEDSC colonies, demonstrating compact colony morphology and AP activity characteristic of pluripotent stem cells. Scale bar, 100 µm. (B) Flow cytometry histograms showing expression of intracellular pluripotency-associated transcription factors POU5F1 (OCT4), SOX2, LIN28A, and NANOG, as well as surface antigens SSEA-4 and SSEA-1, in siPSCs and sEDSCs. Shaded histograms represent marker-positive populations relative to isotype controls, with percentages indicating the proportion of positive cells. (C) Quantitative RT-qPCR analysis of pluripotency-associated gene expression (POU5F1, NANOG, ZFP42, PRDM14, GBX2, LIN28A, SALL4, and DNMT3B) across independent siPSC and sEDSC cell banks. Expression levels were normalized to sheep fibroblasts (sFibs) and to the housekeeping gene HMBS, and relative expression was calculated using the ΔΔCt method. (D) Representative G-band karyotype analyses of siPSCs and sEDSCs demonstrating maintenance of a normal ovine chromosomal complement (2n = 54).

To further compare pluripotent identity at the protein level, expression of established surface and intracellular pluripotency markers was assessed by flow cytometry. sEDSCs predominantly expressed the surface antigen SSEA-4, with comparatively lower levels of SSEA-1, whereas siPSCs displayed the reciprocal pattern, with higher SSEA-1 expression relative to SSEA-4 (Figure 2B). Despite these surface marker differences, both cell types showed comparable intracellular expression of LIN28A. In contrast, the core pluripotency transcription factors POU5F1, SOX2, and NANOG were detected at higher levels in sEDSCs than in siPSCs (Figure 2B), indicating quantitative differences in marker expression rather than absence of pluripotency.

Pluripotency-associated gene expression was further evaluated by quantitative RT-PCR across multiple independently derived cell banks. Both sEDSCs and siPSCs expressed a panel of pluripotency-related genes, including *ZFP42, PRDM14, GBX2, LIN28A, SALL4*, and *DNMT3B* (Figure 2C). Analysis of three independent cell banks per cell type revealed some degree of clonal variability within both sEDSC and siPSC groups; however, overall expression profiles were consistent with pluripotent stem cell identity. Gene expression levels were normalized to the housekeeping gene *HMBS* and expressed relative to sheep fibroblasts, which served as a differentiated somatic reference.

Finally, cytogenetic stability was assessed to confirm suitability of both platforms for downstream applications. Karyotype analysis demonstrated that both siPSCs and sEDSCs maintained a normal ovine chromosome complement (2n = 54), with no gross chromosomal abnormalities detected in the analyzed metaphase spreads (Figure 2D).

### Differentiation potential

To assess functional pluripotency, both siPSCs and sEDSCs were evaluated for their ability to undergo multilineage differentiation *in vitro*. Trilineage differentiation was first examined using an embryoid body–based assay. Following dissociation and aggregation, both siPSCs and sEDSCs efficiently formed embryoid bodies within 3 days of induction (Figure 3A). These structures subsequently attached and underwent further differentiation over the following days. By day 10 of differentiation, expression of lineage-specific markers representing the ectoderm, mesoderm, and endoderm germ layers was detected in all cell lines, confirming trilineage differentiation capacity for both pluripotent stem cell platforms (Figure 3B).

**Figure 3.**
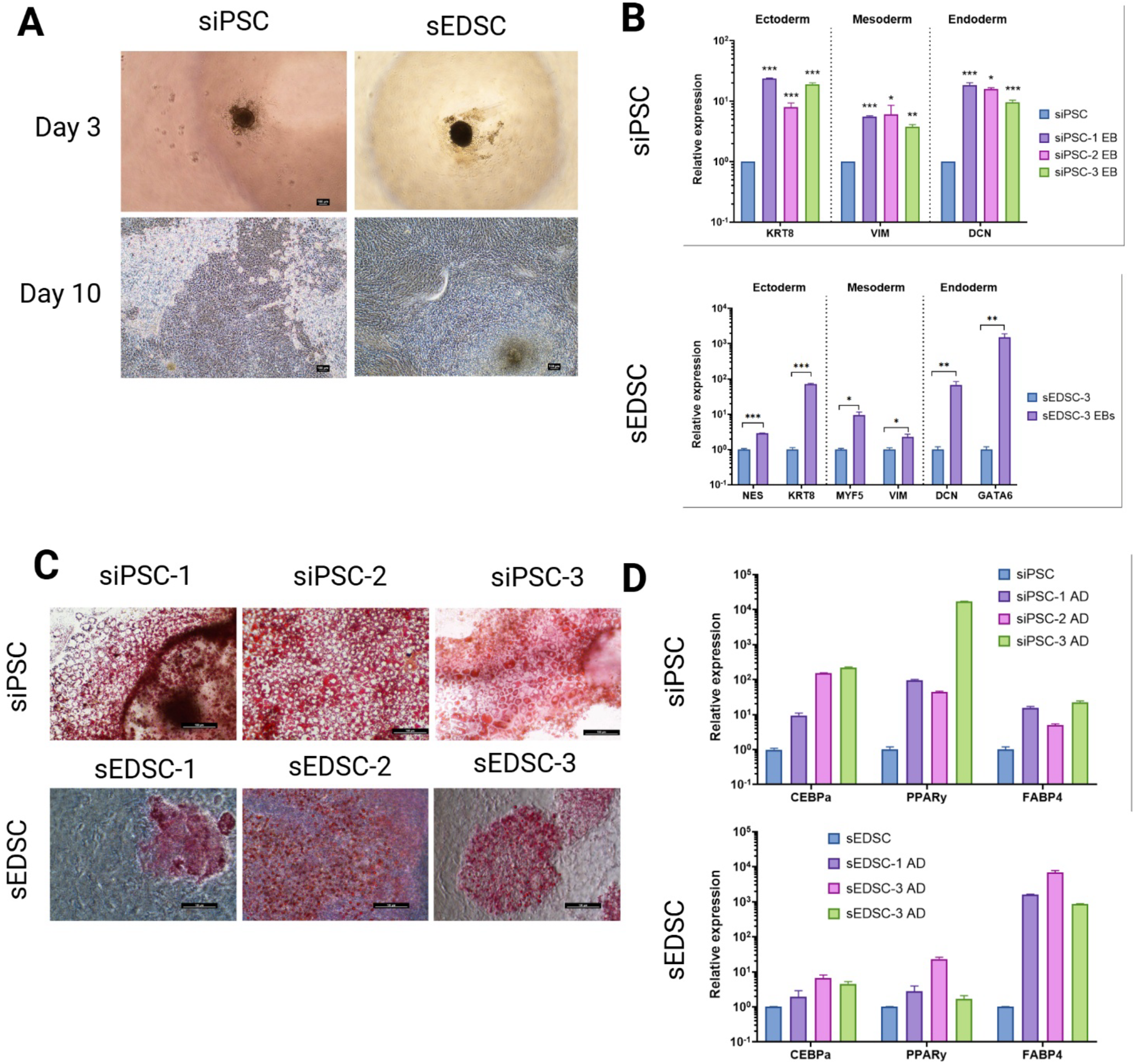
Assessment of multilineage and adipogenic differentiation potential of ovine induced pluripotent stem cells (siPSCs) and embryonic disc-derived stem cells (sEDSCs). (A) Representative phase-contrast images of embryoid body (EB) formation at day 3 of differentiation (suspension culture) and EB outgrowths at day 10 following attachment for siPSCs and sEDSCs. Scale bar, 100 µm. (B) Quantitative RT-qPCR analysis of lineage-associated marker expression following EB-based differentiation. Upper panel: siPSC derivatives showing ectodermal (KRT8), mesodermal (VIM), and endodermal (DCN) marker expression. Lower panel: sEDSC derivatives showing expression of ectodermal (NES, KRT8), mesodermal (MYF5, VIM), and endodermal (DCN, GATA6) markers. (C) Representative bright-field images of Oil Red O staining following adipogenic differentiation of independent siPSC (siPSC-1, siPSC-2, siPSC-3) and sEDSC (sEDSC-1, sEDSC-2, sEDSC-3) lines, demonstrating intracellular lipid accumulation. Scale bars as indicated. (D) Quantitative RT-qPCR analysis of adipogenic marker expression (CEBPα, PPARγ, and FABP4) following adipogenesis.

Given the relevance of mesodermal derivatives for potential downstream applications, we next examined the capacity of siPSCs and sEDSCs to differentiate toward the adipogenic lineage. Under adipogenic differentiation conditions, both cell types showed evidence of adipocyte formation; however, marked differences in differentiation efficiency were observed. siPSCs consistently exhibited robust accumulation of intracellular lipid droplets, as visualized by Oil Red O staining (Figure 3C). In contrast, sEDSCs displayed weaker and less uniform lipid staining under the same induction conditions.

These qualitative observations were supported by transcriptional analysis of adipogenic marker expression. Differentiated siPSCs exhibited higher expression levels of key adipogenic genes, including *FABP4, CEBPα*, and *PPARγ*, compared with differentiated sEDSCs (Figure 3D). This data indicates that while both sEDSCs and siPSCs retain the capacity for multilineage differentiation, their responsiveness to adipogenic induction differs, with siPSCs showing enhanced adipogenic differentiation efficiency under the conditions tested.

### Genetic stability and transcriptomics

To assess transcriptomic identity and genetic stability of the derived pluripotent stem cell platforms, we performed bulk RNA sequencing on siPSCs, sEDSCs, and their corresponding somatic donor cell types (neural cells and fibroblasts). Principal component analysis of global gene expression profiles demonstrated a clear separation between pluripotent stem cells and somatic populations, with siPSCs and sEDSCs clustering closely together and distinctly from neural cells and fibroblasts (Figure 4A). This clustering pattern indicates a shared pluripotent transcriptional signature across both stem cell platforms despite their distinct derivation routes.

**Figure 4.**
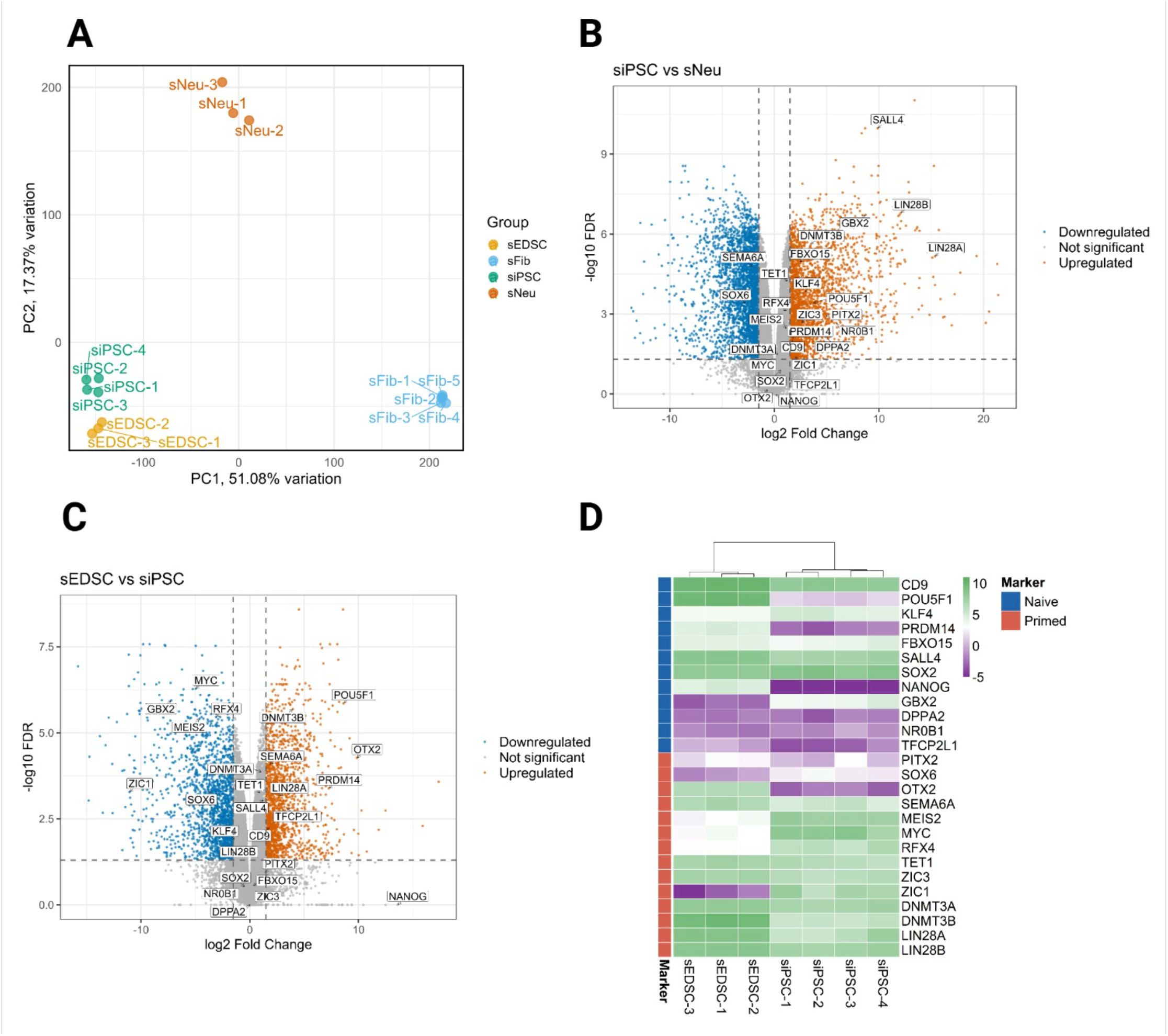
Global transcriptomic analysis of ovine induced pluripotent stem cells (siPSCs), embryonic disc-derived stem cells (sEDSCs), and somatic cell populations. (A) Principal component analysis (PCA) of bulk RNA-sequencing data showing clustering of siPSC and sEDSC samples relative to primary neural cells and fibroblasts, based on global gene expression profiles. Each point represents an individual biological replicate. (B) Volcano plot showing differential gene expression between siPSCs and neural somatic cells. Significantly up-regulated and down-regulated genes are highlighted based on false discovery rate and fold-change thresholds, with selected pluripotency-associated genes indicated. (C) Volcano plot showing global gene differentially expression between sEDSCs and siPSCs under defined statistical thresholds, highlighting quantitative transcriptional differences between the two pluripotent stem cell platforms. (D) Heatmap depicting relative expression patterns of genes associated with naïve- and primed-like pluripotency states in siPSCs and sEDSCs. Gene expression values are scaled across samples, with hierarchical clustering applied to both genes and samples.

Differential expression analysis comparing siPSCs with their neural starting population identified 19,324 genes passing filtering thresholds, of which 5,996 genes were up-regulated in siPSCs and 6,129 genes were up-regulated in neural cells, based on an FDR < 0.05 and an absolute log_2_ fold-change ≥ 1.5 (Figure 4B). The remaining genes did not meet significant thresholds. Genes up-regulated in siPSCs included a range of established pluripotency-associated markers, encompassing factors previously reported in both naïve- and primed-like pluripotent states. This transcriptional shift confirms successful reprogramming of neural cells and acquisition of a pluripotent expression profile distinct from the parental somatic state.

Transcriptomic comparison between sEDSCs and siPSCs revealed a similarly broad set of expressed genes, with 19,324 genes analyzed under the same statistical criteria. Within this set, 4,679 genes were up-regulated in sEDSCs, while 4,305 genes were up-regulated in siPSCs (Figure 4C), with the remainder showing no significant differential expression. This differential expression analysis identified subsets of genes that were differentially regulated between siPSCs and sEDSCs, reflecting quantitative divergence within an otherwise related pluripotent gene expression landscape. Importantly, many genes associated with core pluripotency and pluripotent state regulation did not segregate strongly between siPSCs and sEDSCs at the transcriptome-wide level, consistent with a shared core pluripotent program.”

To further examine pluripotency state characteristics, expression patterns of genes associated with naïve and primed pluripotency were analyzed across both datasets. Both siPSCs and sEDSCs expressed markers commonly linked to naïve- and primed-like pluripotent states, with no clear segregation into a single canonical pluripotency category (Figure 4D). This mixed expression profile is consistent with observations in other livestock pluripotent stem cell systems and supports the conclusion that both ovine pluripotent stem cell platforms occupy closely related, but not identical, pluripotent transcriptional states.

### Long-term stability

To evaluate long-term stability during extended culture, both siPSCs and sEDSCs were assessed across early, mid, and late passages for maintenance of morphology, proliferation characteristics, pluripotency marker expression, and genomic integrity. siPSC cultures were examined from passage 10 through passage 51, while sEDSC cultures were analyzed from passage 21 through passage 38, reflecting sustained expansion of both pluripotent stem cell platforms.

Throughout prolonged passaging, both siPSCs and sEDSCs retained colony morphologies characteristic of pluripotent stem cells and remained positive for AP activity at all analyzed time points (Figure 5A). No overt signs of spontaneous differentiation or loss of colony integrity were observed during continued culture, indicating stable maintenance of the undifferentiated state.

**Figure 5.**
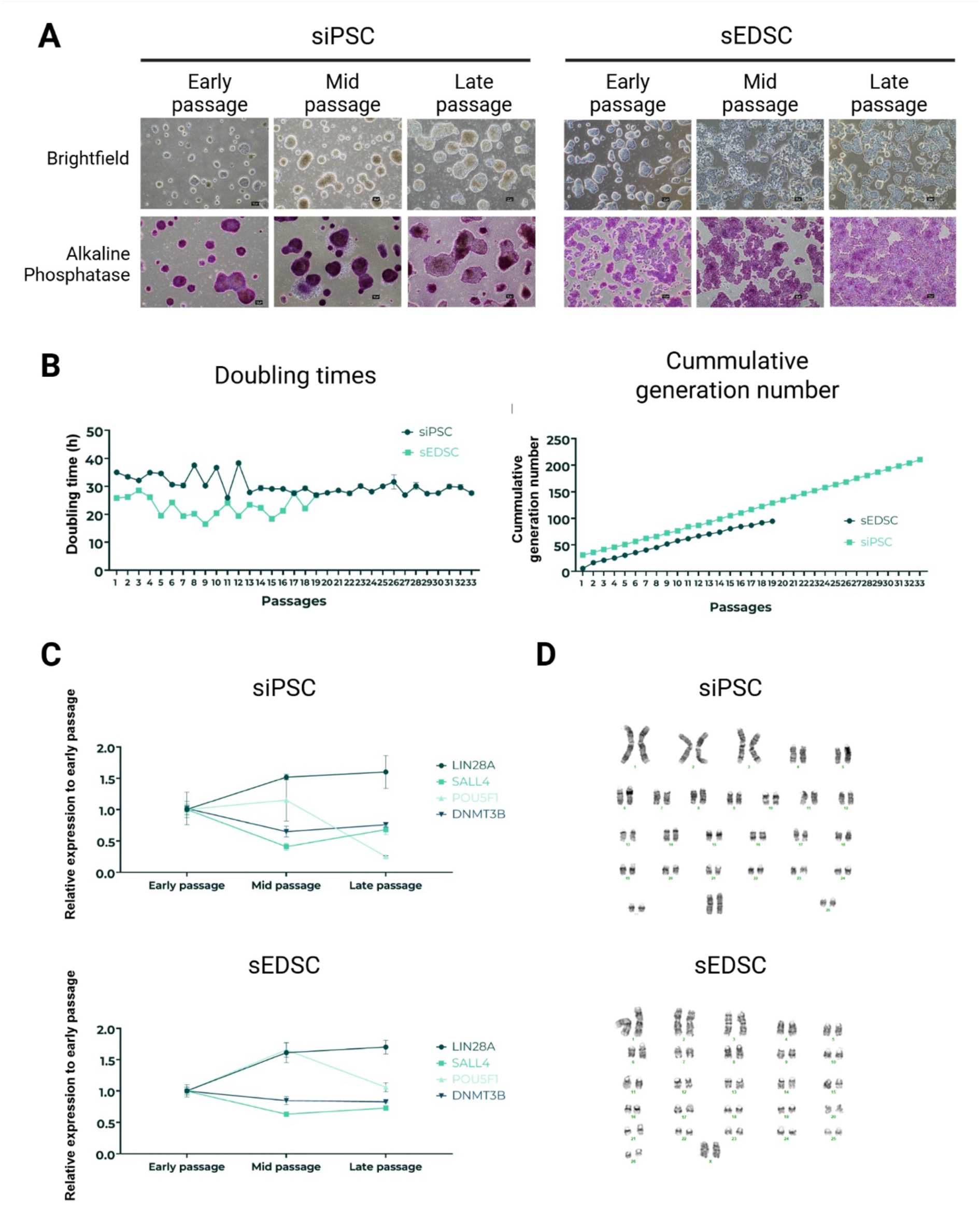
Evaluation of morphological, proliferative, molecular, and chromosomal stability of ovine pluripotent stem cells across serial passaging. (A) Representative phase-contrast images (top row) and alkaline phosphatase staining (bottom row) of siPSC and sEDSC cultures at early, mid, and late passages. Scale bar, 100 µm. (B) Growth kinetics of siPSCs and sEDSCs over serial passaging, shown as population doubling times (left) and cumulative population doublings (right), calculated from triplicate cell counts at each passage. (C) Quantitative RT-qPCR analysis of pluripotency-associated gene expression (LIN28A, SALL4, POU5F1, and DNMT3B) in siPSCs (upper panel) and sEDSCs (lower panel) at early, mid, and late passages. Expression levels were normalized to early-passage cultures and to the housekeeping gene HMBS. (D) Representative G-band karyotype analyses of late-passage siPSC and sEDSC cultures demonstrating preservation of a normal ovine chromosomal complement (2n = 54) following long-term culture.

Growth kinetics were quantified to assess proliferative behavior over time. siPSCs exhibited an average population doubling time of 30.7 ± 2.0 hours, corresponding to a cumulative population doubling level of 210 across the analyzed passage range. In comparison, sEDSCs displayed a shorter average doubling time of 22.7 ± 3.4 hours, with a cumulative population doubling level of 94 (Figure 5B). These data demonstrate consistent, long-term proliferative capacity for both cell types, while highlighting quantitative differences in growth behavior between embryonic-derived and induced pluripotent stem cells.

Maintenance of pluripotent identity during long-term culture was further evaluated by quantitative RT-PCR analysis of core pluripotency-associated genes. Expression levels of *LIN28A, SALL4, POU5F1*, and *DNMT3B* remained relatively stable across early, mid, and late passages for both siPSCs and sEDSCs, with relative expression values ranging between 0.4 and 1.8 when normalized to early-passage cultures (Figure 5C). These results indicate sustained expression of key pluripotency markers during extended expansion.

Finally, chromosomal stability was assessed at the end of the culture period for each cell type. Karyotype analysis of late-passage siPSC and sEDSC cultures revealed a normal ovine chromosome complement (2n = 54), with no gross chromosomal abnormalities detected in the analyzed metaphase spreads (Figure 5D).

Together, these data demonstrate that both ovine siPSCs and sEDSCs maintain morphological, molecular, proliferative, and genetic stability during long-term *in vitro* expansion, supporting their suitability as robust pluripotent stem cell platforms for downstream applications.

## Discussion

Pluripotency is best understood not as a fixed cellular identity but as a transient and context-dependent regulatory state that emerges during early development and can be artificially stabilized *in vitro* under defined conditions (1,28,29) In this study, we exploit two fundamentally distinct routes to the acquisition of pluripotency in sheep—direct derivation from the embryonic disc and transcription factor-mediated reprogramming of somatic cells—to examine how developmental origin, route to pluripotency, and stabilizing culture conditions shape pluripotent state identity, regulatory architecture, and downstream functional behavior. By integrating morphological, molecular, transcriptional, and functional analyses, our data provide insight into the extent to which embryo-derived and reprogrammed pluripotent stem cells share a common core of pluripotency while occupying distinct regulatory configurations with measurable consequences for differentiation responsiveness and translational utility.

Sheep embryonic disc-derived stem cells (sEDSCs) are captured directly from embryonic pluripotent tissue and therefore represent a pluripotent state that is developmentally continuous with the lineage trajectory of the inner cell mass and post-implantation epiblast. In mammals, pluripotency *in vivo* is highly transient and progresses through a series of molecularly distinct configurations as the epiblast transitions from naïve to formative and primed states. Pluripotency is maintained by a metastable transcription factor network centered on OCT4, SOX2, and NANOG, whose stability depends on the integration of intrinsic transcriptional feedback with extrinsic signaling cues rather than on irreversible stem cell identity (28–30). Importantly, this framework predicts that pluripotent cells captured at different developmental windows, or stabilized by different signaling environments, may exhibit distinct phenotypic and functional properties without loss of pluripotent competence.

Consistent with this view, sEDSCs in the present study exhibited stable colony morphology, robust alkaline phosphatase activity, and relatively elevated expression of core pluripotency-associated transcription factors. Their flatter colony architecture is reminiscent of formative or primed-like pluripotent states rather than a naïve mouse embryonic stem cell like state, in which dome-shaped, three-dimensional colonies are typically observed. While such morphological correlations have historically been used to infer pluripotent state in mouse and human systems (31,32), direct extrapolation across species is problematic. Indeed, recent embryo-aligned transcriptomic analyses in sheep demonstrate that pluripotency most closely associated with the embryonic disc does not map cleanly onto canonical naïve or primed states defined in rodents, instead occupying intermediate or formative configurations shaped by species-specific developmental timing and signaling dependencies (20,33). The properties of sEDSCs observed here are therefore consistent with a pluripotent state that remains tightly coupled to endogenous developmental programs rather than with diminished pluripotent potential.

In contrast to embryo-derived pluripotency, induced pluripotent stem cells are generated through an explicitly artificial process in which ectopic expression of lineage-defining transcription factors overrides a pre-existing somatic gene regulatory network. Although reprogramming collapses much of somatic identity and re-establishes expression of core pluripotency regulators, it does not simply recreate embryonic pluripotency. iPSCs represent stabilized endpoints within a landscape of transcriptionally induced states whose properties reflect both the starting cell type and the imposed reprogramming and culture environment (1,29,30). Accordingly, siPSCs in this study displayed hallmark features of pluripotency, including trilineage differentiation capacity and stable self-renewal, yet differed from sEDSCs in colony architecture, surface antigen expression, and quantitative transcription factor abundance.

Variation in pluripotency marker expression has been widely reported in livestock pluripotent stem cells and highlights the limitations of defining pluripotent state based on single markers or antibody panels. Early studies reported inconsistent detection of SSEA-1, SSEA-4, and NANOG across ovine ESC-like cells and iPSCs, differences that have been attributed to species-specific regulatory networks, technical variability, and limited reagent cross-reactivity rather than to absence of pluripotency itself (11,13,33–35). Within the transcriptional regulation framework, such variability is expected as pluripotency is defined by the topology and stability of a distributed gene regulatory network, not by absolute expression of individual canonical markers. The marker expression patterns observed in both sEDSCs and siPSCs in this study are therefore best interpreted as reflecting quantitative differences in pluripotent state configuration rather than divergent pluripotency status.

Transcriptomic profiling reinforces this interpretation. RNA sequencing confirmed that siPSCs undergo extensive transcriptional reprogramming relative to their neural progenitor origin, yet global comparisons revealed that siPSCs and sEDSCs occupy closely related pluripotent transcriptional spaces distinct from somatic populations. Notably, neither cell type segregated cleanly into naïve or primed categories based on established marker gene sets. This finding aligns with recent embryo-based transcriptomic studies demonstrating that ovine pluripotency exists along a continuum of regulatory states rather than as discrete categories and further emphasizes the importance of species-specific frameworks when interpreting pluripotent identity in large mammals (20,33). Accordingly, transcriptomic similarity between siPSCs and sEDSCs should be inferred from their shared global expression structure and clustering relative to somatic cells, rather than from the absolute number of globally differentially expressed genes identified.

At a mechanistic level, differences between sEDSCs and siPSCs likely arise from how core pluripotency transcription factors integrate with signaling pathways such as FGF/ERK, TGF-β/activin, WNT/β-catenin, and JAK/STAT. Small shifts in pathway balance can stabilize alternative pluripotent attractor states with distinct lineage biases and differentiation responsiveness. In embryo-derived cells, this balance is established progressively through development; in reprogrammed cells, it is imposed abruptly and may stabilize in configurations that differ subtly but meaningfully from embryonic counterparts. Such differences are increasingly recognized as central to understanding why iPSCs can display altered differentiation propensities despite satisfying formal criteria for pluripotency.

Functional consequences of these regulatory distinctions were evident during directed differentiation. Although both sEDSCs and siPSCs demonstrated trilineage differentiation capacity, siPSCs exhibited a stronger adipogenic response under the conditions tested. This enhanced differentiation may reflect epigenetic memory associated with the neural starting population, altered chromatin accessibility at mesodermal loci, or heightened sensitivity to exogenous differentiation cues, phenomena that have been documented in multiple iPSC systems (35–37). Conversely, the more restrained adipogenic response of sEDSCs may reflect a pluripotent state in which lineage priming is more tightly regulated, consistent with their closer developmental relationship to embryonic disc epiblast.

These observations have direct implications for the application of pluripotent stem cells in cellular agriculture and cultivated meat production. While pluripotent stem cells offer unparalleled scalability and developmental versatility, not all pluripotent states are equally suited for industrial differentiation pipelines. siPSCs offer clear advantages in terms of sourcing flexibility, donor specificity, and renewability, and their enhanced responsiveness to adipogenic differentiation may be advantageous for generating fat components that contribute to flavor and texture in cultivated meat products. However, their artificial origin necessitates careful evaluation of long-term stability, reproducibility, and regulatory acceptability. sEDSCs, by contrast, provide a developmentally grounded reference state that may offer greater predictability and regulatory coherence but are constrained by ethical, logistical, and scalability considerations inherent to embryo-derived systems.

Taken together, our findings reinforce the concept that pluripotency encompasses a spectrum of regulatory states shaped by developmental history and technological intervention. By directly comparing embryo-derived and reprogrammed pluripotent stem cells in sheep within a unified experimental framework, this study situates ovine pluripotency within a broader conceptual landscape defined by transcriptional regulation, signal integration, and functional competence. Such understanding is essential for rational selection and refinement of pluripotent cell substrates for both fundamental developmental biology and emerging applications in livestock biotechnology and cultivated meat production, where functional output, scalability, and regulatory alignment are as critical as pluripotent identity itself.

## Declarations

### Funding

This work was funded by Roslin Technologies, UK Research and Innovation (10014258) and Scottish Enterprise SMART award.

### Author Contributions

ASA, MED, WA and SH performed cell derivation, culture, and *in vitro* differentiation experiments. MP, and SW conducted transcriptomic analyses and data processing. DR and PJM contributed to experimental design, data interpretation, and project supervision. ASA drafted the manuscript. DR and PJM critically revised the manuscript. All authors reviewed and approved the final version of the manuscript.

### Conflict of Interest

The authors declare that the research was conducted in the absence of any commercial or financial relationships that could be construed as a potential conflict of interest. All authors are employees of Roslin Technologies Ltd.; however, this affiliation did not influence the study design, data collection, analysis, interpretation, or the decision to publish the results.

### Data Availability Statement

All data supporting the conclusions of this study are included within the article. The RNA-sequencing datasets generated during this study are subject to commercial restrictions but are available from Roslin Technologies Ltd. servers and can be accessed upon reasonable request from the corresponding author. All proprietary medias are available from appropriate suppliers and used according to manufactures instructions. All other data supporting the findings of this study, including quantitative PCR results, flow cytometry analyses, and figure source data, are included within the article and/or provided as supplementary material.

### Ethics and tissue sourcing

All ovine biological materials used in this study were obtained in accordance with institutional, national UK Animals (Scientific Procedures) Act, and international guidelines for the use of animal-derived tissues in research. Sheep embryos and postnatal tissues were sourced from a licensed commercial abattoir as by-products of routine agricultural practices. No animals were euthanized specifically for the purpose of this study, and no formal ethical approval was required under local institutional and national regulations.

## Acknowledgments

The authors thank staff at Roslin Technologies for technical support, cell banking and quality control assays. We would like to thank the staff at ABreeds for support with embryo work especially Geraint Thomas and Fenranda Aidar as well as the staff at Wishart Abattoir for access to primary tissue

## Significance Statement

Pluripotent stem cells from livestock species are emerging as valuable tools for studying early development and enabling applications such as cellular agriculture, yet their biological properties remain incompletely defined. This study provides a direct comparison of ovine induced pluripotent stem cells and embryonic disc-derived stem cells generated under aligned conditions. We demonstrate that both platforms share core features of pluripotency, including transcriptional identity, differentiation capacity, and long-term stability, while exhibiting distinct molecular profiles and functional differences in lineage-specific differentiation. These findings advance understanding of pluripotent state diversity in large mammals and establish a framework for selecting and optimizing stem cell platforms for research and biotechnological applications.

## Notes

### Competing Interest Statement

The authors have declared no competing interest.

### Summary of Updates

The author list is now correctly Reordered

